# MultiCapsNet: a interpretable deep learning classifier integrate data from multiple sources

**DOI:** 10.1101/570507

**Authors:** Lifei Wang, Xuexia Miao, Jiang Zhang, Jun Cai

**Author notes:** Correspondence to: Jiang Zhang, Jun Cai.

## Abstract

Recent advances in experimental biology have generated huge amount of data. Due to differences present in detection targets and detection mechanisms, the produced data comes with different formats and lengths. There is an urgent call for computational methods to integrate these diverse data. Deep learning model is an ideal tool to cope with complex datasets, but its inherent ‘black box’ nature needs more interpretability. Here, we present MultiCapsNet, a deep learning model built on CapsNet and scCapsNet. The MultiCapsNet model possesses the merits of both easier data integration and higher model interpretability. In the first example, we use the labeled variant call dataset, which is originally used to test the models for automating somatic variant refinement. We divide the 71 features listed in the dataset into eight groups according to data source and data property. Then, the data from those eight groups with different formats and lengths are integrated by our MultiCapsNet to predict the labels associated with each variant call. The performance of our MultiCapsNet matches the previous deep learning model well, given much less parameters than those needed by the previous model. After training, the MultiCapsNet model provides importance scores for each data source directly, while the previous deep learning model needs an extra importance determination step to do so. Despite that our MultiCapsNet model is substantially different from the previous deep learning model and the source importance measuring methods are also different, the importance score correlation between these two models is very high. In the second example, the prior knowledge, including information for protein-protein interactions and protein-DNA interactions, is used to determine the structure of MultiCapsNet model. The single cell RNA sequence data are decoupled into multiple parts according to the structure of MultiCapsNet model that has been integrated with prior knowledge, with each part represents genes influenced by a transcription factor or involved in a protein-protein interaction network and then could be viewed as a data source. The MultiCapsNet model could classify cells with high accuracy as well as reveal the contribution of each data source for cell type recognition. The high ranked contributors are often relevant to the contributed cell type.

## Introduction

Recent advances in experimental biology have generated huge amount of data. More detectable biological targets and various new measuring methods produce data at an unprecedented rate. For example, microwell-seq, a single-cell RNA sequencing technique, have been used to analyze the transcriptome of more than 400,000 mouse single cells, which are covering all of the major mouse organs [1]; single-cell bisulfite sequencing (scBS-seq) is designed to measure DNA methylation at the single-cell level [2]; Mass-spectrometry-based technologies could explore the composition, structure, function and control of the proteome [3]. Furthermore, huge complex datasets are generated by large projects such as ‘The Cancer Genome Atlas’ (TCGA) [4] and ‘Encyclopedia of DNA Elements’ (ENCODE) [5], which are built up through the community collaboration. There is an urgent call for next-generation methods to handle large, heterogeneous, complex datasets [6].

As a promising data processing method, deep learning methods have been applied to process biological data [6–8]. Various deep learning models could deal with various input data with different types and formats. For example, RNA sequence data, which are real-value vector, could be processed by simple feed forward neural network which is a constituent of more complex model such as auto-encoder (AE) [9, 10], variational auto-encoder (VAE) [11] and Generative adversarial network (GAN) [12]. Sequence information, which is coded by ATCG, could be converted into real value vector by deep learning model using convolution neural networks (CNN) after model training [8]. Furthermore, deep learning model could integrate data with different types and formats. For example, DeepCpG utilizes both DNA sequence patterns and neighboring methylation states for predicting single-cell methylation states and modeling the sources of DNA methylation variability[13]. But the deep learning methods, which usually operate as a ‘black box’, are hard to interpret[14]. There have been substantial efforts to increase the interpretability of deep learning model. The prior biological information, such as influence between transcriptional factors (TF) and regulated genes, could specify connections between neurons in the neural networks in order to associate the inter node (neuron) in the neural networks with transcriptional factors and thereby ease the difficulty of interpreting models[9]. New probabilistic generative model with more interpretability, such as variational inference neural networks, are applied to single cell transcriptome data for dimension reduction [11].

The CapsNet is a deep learning model which is used in the task of digit recognition and exhibits more interpretability [15].In the realm of biology, the CapsNet model have been applied for protein structure classification and prediction [16, 17] and is ripe for application in network biology and disease biology with data from multi-omics dataset [6]. In our previous work, we propose a modified CapsNet model suitable for single-cell RNA sequencing (scRNA-seq) data, which is called scCapsNet [18]. The scCapsNet is capable of extracting features from scRNA-seq data, and then uses those extracted features for cell type classification with contribution of each feature to type recognition as co-product.

Here, we present MultiCapsNet, a deep learning model builds on CapsNet and scCapsNet. The MultiCapsNet model is supposed to deal with data from multiple sources with different formats and lengths, and meanwhile give the importance scores of each data source for prediction after training. In the first example, we use the labeled variant call dataset, which is originally used to test the models for automating somatic variant refinement [19]. We group the 71 features listed in the dataset into eight groups according to data source and data property, and then view the features in one group as a whole to train the MultiCapsNet model. The performance of our MultiCapsNet matches the previous deep learning model well, given much less parameters than those needed by the previous model. After training, our MultiCapsNet model provides the importance score for each data source, while the previous deep learning model needs an extra importance determination step through shuffling individual features. Despite that our MultiCapsNet model is substantially different from the previous deep learning model and the source importance measuring methods are also different, the importance score correlation between these two models is very high.

In second example, we demonstrate how to integrate prior knowledge, which could be viewed as data sources, and scRNA-seq data into MultiCapsNet model. The protein-protein interaction (PPI) information stored in BIOGRID [20] and HPRD [21], and protein-DNA interaction (PDI) information from [22], are used as prior knowledge to specify network connections like previous work [9]. In this example, the structures of the first part of MultiCapsNet model, which are the connections between inputs and primary capsules, are determined by the PPI and PDI information. The consequence of these specified structures is that each primary capsule is labeled either as transcription factor (TF) or PPI cluster nodes (PPI), and input for each primary capsule could be viewed as a data source. We use data from mouse scRNA-seq dataset [1] to train this MultiCapsNet model. The accuracy of classification is very high. After training, the MultiCapsNet model reveals how each primary capsule, which is labeled either as transcription factor (TF) or PPI cluster nodes (PPI) and receives data from a data source, contributes to cell type classification. The top ranked contributors for one specific cell type often associate with this cell type.

## Methods

### Datasets and data preprocessing

The data used to test the MultiCapsNet model comes from previous work [19]. The dataset, which contains more than 41000 samples, is assembled to train models for automating somatic variant refinement. Each sample in the dataset is manually tagged by reviewer as one of the four labels: ‘somatic’, ‘ambiguous’, ‘germline’ and ‘fail’, which represent the confidence of a variant call by upstream somatic variant caller. As previous work, we merge variant calls labeled as ‘germline’ and ‘fail’ into one class called ‘fail’. There are 71 features that associate with each sample, including cancer types, reviewers, tumor read depth, normal read depth, and so on. We divide the 71 features into eight groups according to data source and data properties (Supplementary Table 1). Group one contains nine cancer types and is coded as one-hot encoding. We label group one as ‘Disease’ because it indicate the disease that each variant call belong to. Group two contains four reviewers and is coded as one-hot encoding. We label group one as ‘Reviewer’. Group three contains information of ‘normal VAF’, ‘normal depth’, ‘normal other bases count’ and is labeled as ‘Normal_pro’, short for ‘Normal properties’. Group four contain thirteen features describe reference reads in normal, including base quality, mapping quality, numbers of mismatches, numbers of minus and plus strand, and so on. We label group four as ‘Normal_ref’. Group five contain thirteen features extracted from variant reads in normal, also including base quality, mapping quality, numbers of mismatches, numbers of minus and plus strand, and so on. We label group five as ‘Normal_var’. The last three groups contain features draw from tumor instead of normal in previous three groups. As labeling of Group three, four and five, we label group six, seven and eight as ‘Tumor_pro’, ‘Tumor_ref’ and ‘Tumor_var’ respectively.

The mouse single cell RNA sequence data, which are measured by microwell-seq, come from [1]. We download scRNA-seq data and annotation information though the link that the authors provide. Then we use the annotation information to select portion of data from whole datasets. The cell types we choose include ‘Cartilage cell’, ‘Neuron’, ‘AT2 Cell’, ‘Stomach cell’, ‘Muscle cell’, ‘Dendritic cell’, and ‘Cumulus cell’. In order to fit the model structure, we only use the genes selected from [9], and set the default value to zero in case that the gene is not comprised in [1].

### MultiCapsNet model

We adapt MultiCapsNet model from previous scCapsNet model. The architecture of MultiCapsNet model is shown in Fig. 1. The model contains two parts, one of which is inputs standardization part and the other is capsule network part. In the scCapsNet, several parallel fully connected neural networks using Rectified Linear Unit (ReLU) activation function are used to extract features from inputs. Each neural network with same structure uses the same input sources, which is the scRNA-seq data. In the MultiCapsNet model, there are multiple input sources instead of one input source in scCapsNet model. For example, in the somatic variant refinement task, we use eight input sources, which are corresponding to eight groups mentioned on the previous section, to classify the variant call. Data from different input source may have different format and length. In the first part of MultiCapsNet model, each input source connects to a fully connected neural networks using Rectified Linear Unit (ReLU) as activation function. The outputs of those neural networks are vectors with equal length. So this part of MultiCapsNet model converts the inputs with different data format and length into vectors with same length and so is called input standardization part. The outputs vectors could be viewed as ‘primary capsule’ in the original CapsNet. Next, the standardized information stored in the primary capsules would be delivered to the final layer capsules by ‘dynamic routing’. The capsules in the final layer, which correspond to labels of variant calls, is called ‘label capsule’. The dynamic routing operation with three iterations occurs between the primary capsule layer and the label capsule layer. In capsule model, the length of the final layer capsule represents the probability that an entity represented by the capsule exists. In our variant call classification task, the length of the final layer label capsule represents the probability that one variant call is either ‘ambiguous’,’ fail’ or ‘somatic’. To evaluate the model performance, we use the AUC (area under the curve) score and the reliability diagram as previous [19].

**Fig. 1:**
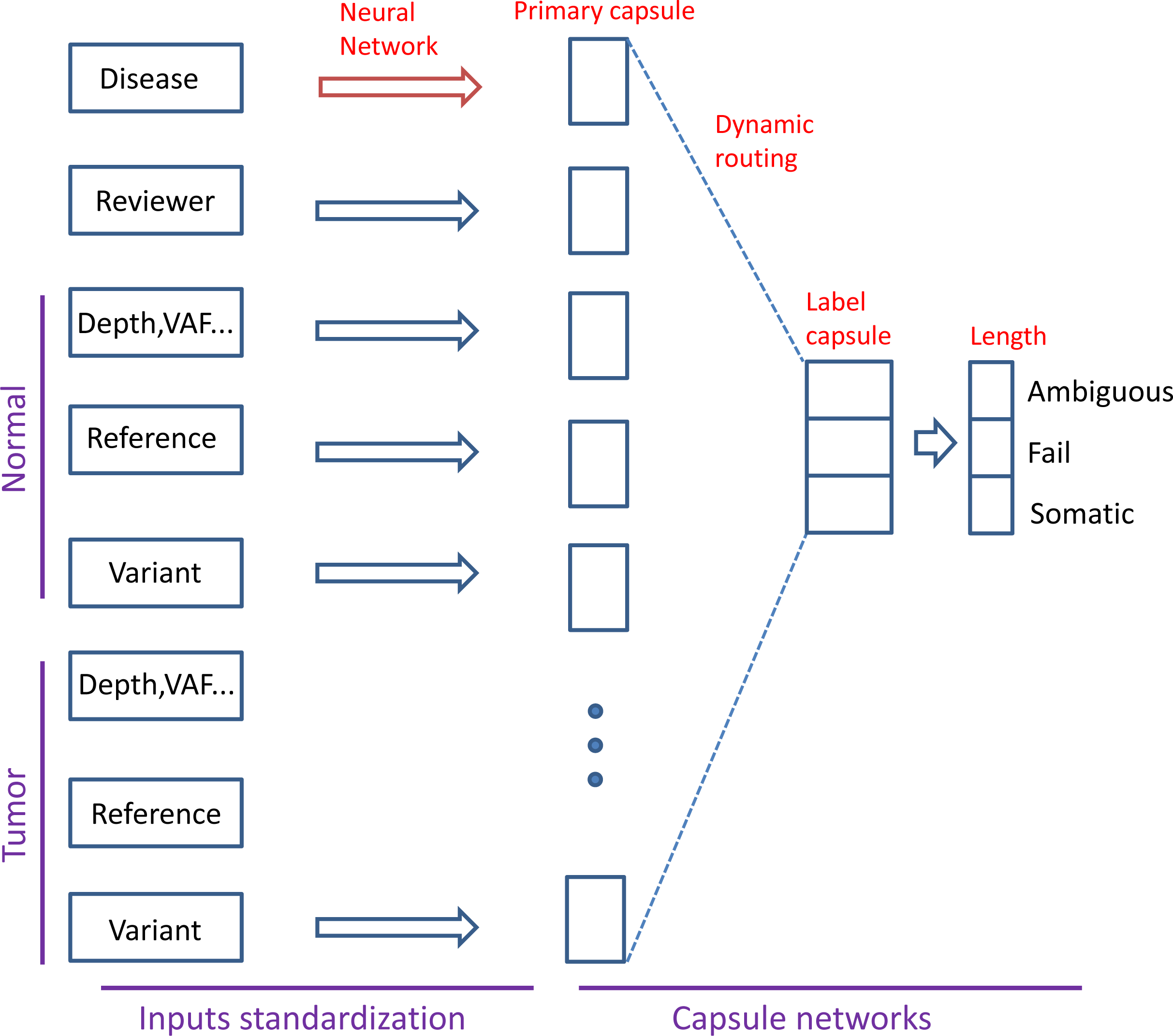
Architecture of MultiCapsNet with two layers. First layer consists of eight parallel fully connected neural networks corresponding to eight data sources (groups). The outputs of the neural network have equal lengths and could be viewed as primary capsules. The second layer is directly adopted from CapsNet for classification. Information flows from primary capsules to final layer label capsules. Then the length of each label capsule represents the probability of input data belonging to the corresponding classification classes.

### MultiCapsNet model that integrates prior knowledge

We use the same method described previously [9] to integrates prior knowledge into model. In short, protein-protein interactions (PPI) information store in BIOGRID[20] and HPRD[21], and protein-DNA interactions (PDI) comes from [22], are used as prior knowledge to specify network connections [9]. In our MultiCapsNet model, each primary capsule is labeled either as transcription factor (TF) or PPI cluster nodes (PPI), and the connections between input nodes and primary capsule are designated by prior knowledge (Fig. 5). For example, if the primary capsule is labeled as TF x, then only the genes (input nodes) that are affected by this TF x, which indicates by PDI, would connect to this primary capsule (TF). Although there is only one input source, which is the scRNA-seq data, the integration of the prior knowledge could decouple this input source into multiple parts, with each part connects to one primary capsule. So, we also view single input source integrated with prior knowledge as inputs from multiple sources and use the networks specify by prior knowledge as first part of MultiCapsNet model. All neural networks in the first part of MultiCapsNet model use tanh as activation function. The second part is capsule networks for cell type classification. The final layer capsule is called ‘Type capsule’.

### Data source importance

The detailed description for dynamic routing could be found in the original CapsNet paper[15]. In MultiCapsNet model, the primary capsules that store the standardized information are first multiplied by a weight matrix to produce ‘prediction vector’. The dynamic routing process would calculate the ‘coupling coefficients’ between each primary capsule’s prediction vector and all final layer capsules. So the total number of coupling coefficients is the product of the number of primary capsules multiplied by the number of final layer capsules. The coupling coefficients of one primary capsule would sum up to one. After dynamic routing, the final layer capsule is actually a weighted sum of prediction vectors of all primary capsules. The weights are the coupling coefficients and the magnitude of those coefficients which indicates the contribution of each primary capsule contributes to the model prediction.

## Results

### Model performance

The dataset we used is the labeled variant call dataset originally used to test the models for automating somatic variant refinement. The dataset contains more than 41000 variant calls, labeled as one of ‘somatic’, ‘ambiguous’, ‘germline’, and ‘fail’. Variant calls tagged as ‘germline’ and ‘fail’ are merged into one class called ‘fail’ as previous for model training [19]. We divide the 71 features associated with each variant call into eight groups according to data source and data property, and then view the features in one group as a whole to train the MultiCapsNet model. The features and their grouping could be found in Supplementary table 1. The eight groups are named as ‘Disease’, ‘Reviewer’, ‘Normal_pro’, ‘Normal_ref’, ‘Normal_var’, ‘Tumor_pro’, ‘Tumor_ref’, and ‘Tumor_var’ respectively. The dataset is randomly divided into training set and validation set with a ratio of 9:1. Our MultiCapsNet model performs well on variant call classification (Fig. 2). The results show that the MultiCapsNet model attains AUC (area under the curve) of 0.94, 0.99 and 0.97 in classification classes of ‘ambiguous’, ‘fail’ and ‘somatic’, respectively (Fig. 2A). These AUC scores are slightly higher than those correspondingly attained by the previous deep learning model (feed forward network with multiple layers) [19], and the parameters of the MultiCapsNet model (700) are less than the previous deep learning model (over 1000). As previous, the reliability diagram is used to evaluate whether the model outputs could be interpreted as a well-scaled probability [19]. The MultiCapsNet model gets Pearson correlation coefficients of 0.96, slightly lower than that yielded by the feed forward neural network (Fig. 2B).

**Fig. 2:**
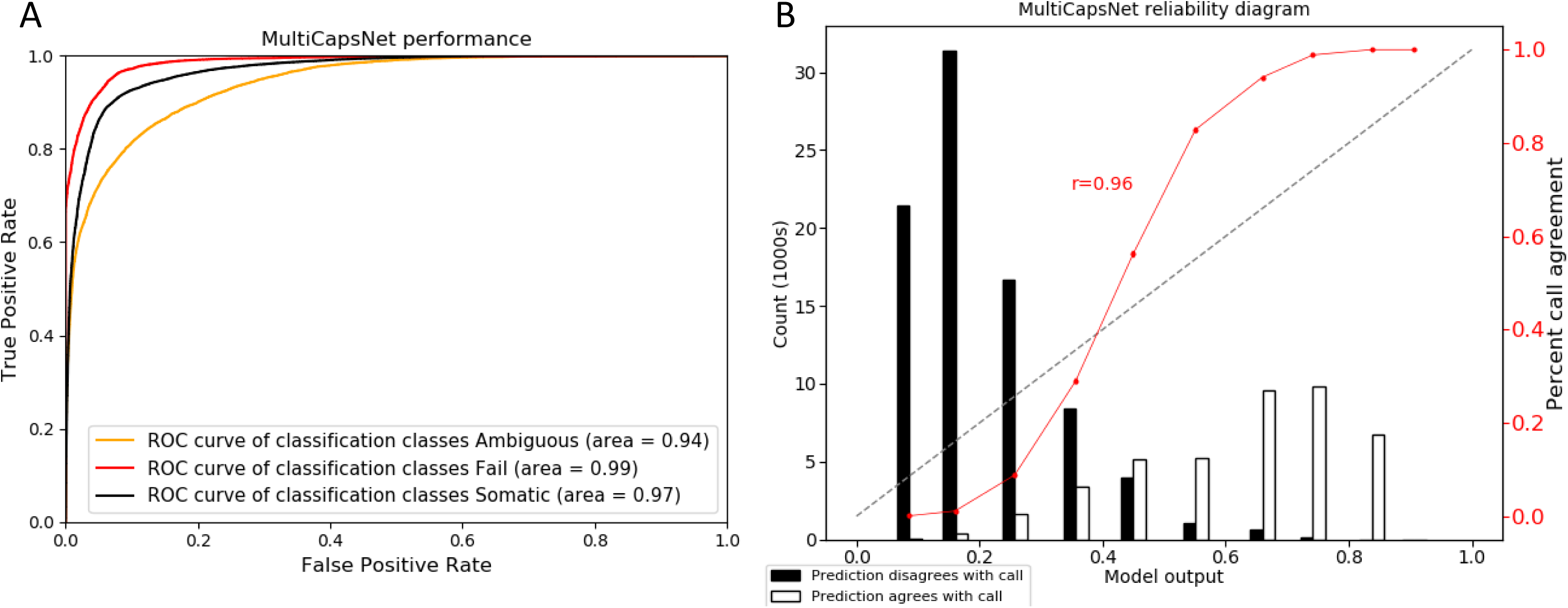
The performance of MultiCapsNet model. **A**. MultiCapsNet model achieves very high classification performances in all three classification classes, which are measured via ROC AUC. **B**. Reliability diagrams measure how the model outputs could be interpreted as a well-scaled probability. The diagonal line indicates a perfectly scaled probabilistic prediction. The red points display the ratio of predictions that agree with the labels, represented by the white bars, to the total number of predictions for a given bin, which is the sum of white bar and black bar. Pearson correlation coefficient comparing red points to the diagonal line is calculated to assess the output of the model.

### Data source importance

In our previous works on cell type classification based on scRNA-seq data, we demonstrate that the coupling coefficients in the trained scCapsNet model could indicate how the extracted features which are stored in primary capsules contribute to cell type classification[18]. In the MultiCapsNet model, the coupling coefficients of the variant calls with the same known variant call label are grouped together and the average coupling coefficients for one specific variant call label are calculated. As previously, the heatmaps visualizations of those average coupling coefficients are plotted (Supplementary Fig. 1A). They show that for input variant call with one specific label, primary capsules with the highest activity contribute to this specific label classification. We also plot the overall heatmap, which incorporates the average coupling coefficients with label capsules corresponding to specific variant call labels from all variant calls (Supplementary Fig. 1B). In scCapsNet model trained on scRNA-seq data, we demonstrate overall heatmap could be viewed as how the primary capsules contribute to the cell type recognition. In MultiCapsNet model, we want to investigate which data sources (eight groups) are important for variant call label classification. So, we add the values for each primary capsules together in the overall heatmap, and use the summed value to measure the contribution of the primary capsules to the variant call type classification, which could be viewed as the data sources importance in MultiCapsNet model (Fig. 3). In the previous deep learning model, the feature importance is measured by average change in the AUC after randomly shuffling individual features. Based on the features grouping step, we add the feature importance values that belong to the same group together, and view those values as data sources (each group) importance in previous deep learning model (Fig. 3). Then, we calculate the correlation between the data source importance values obtained by our MultiCapsNet model and those provided by previous deep learning model. Despite that our MultiCapsNet model is substantially different from the previous deep learning model and the source importance measuring methods are also different, their importance correlation are very high (r^2^ = 0.841) (Fig. 3). Both models indicate the group ‘Tumor_var’ is very important for variant call classification. We also run MultiCapsNet model several time with different dataset division, which divide the dataset into training set and validation set, in order to test the model consistency. As plot show in Fig. 4, MultiCapsNet model performs relatively stable among different rounds, with median AUC of 0.965, median MultiCapsNet model importance score correlation of 0.956 between pair of any two rounds, and median importance score correlation of 0.841 between MultiCapsNet model and previous deep learning model.

**Fig. 3:**
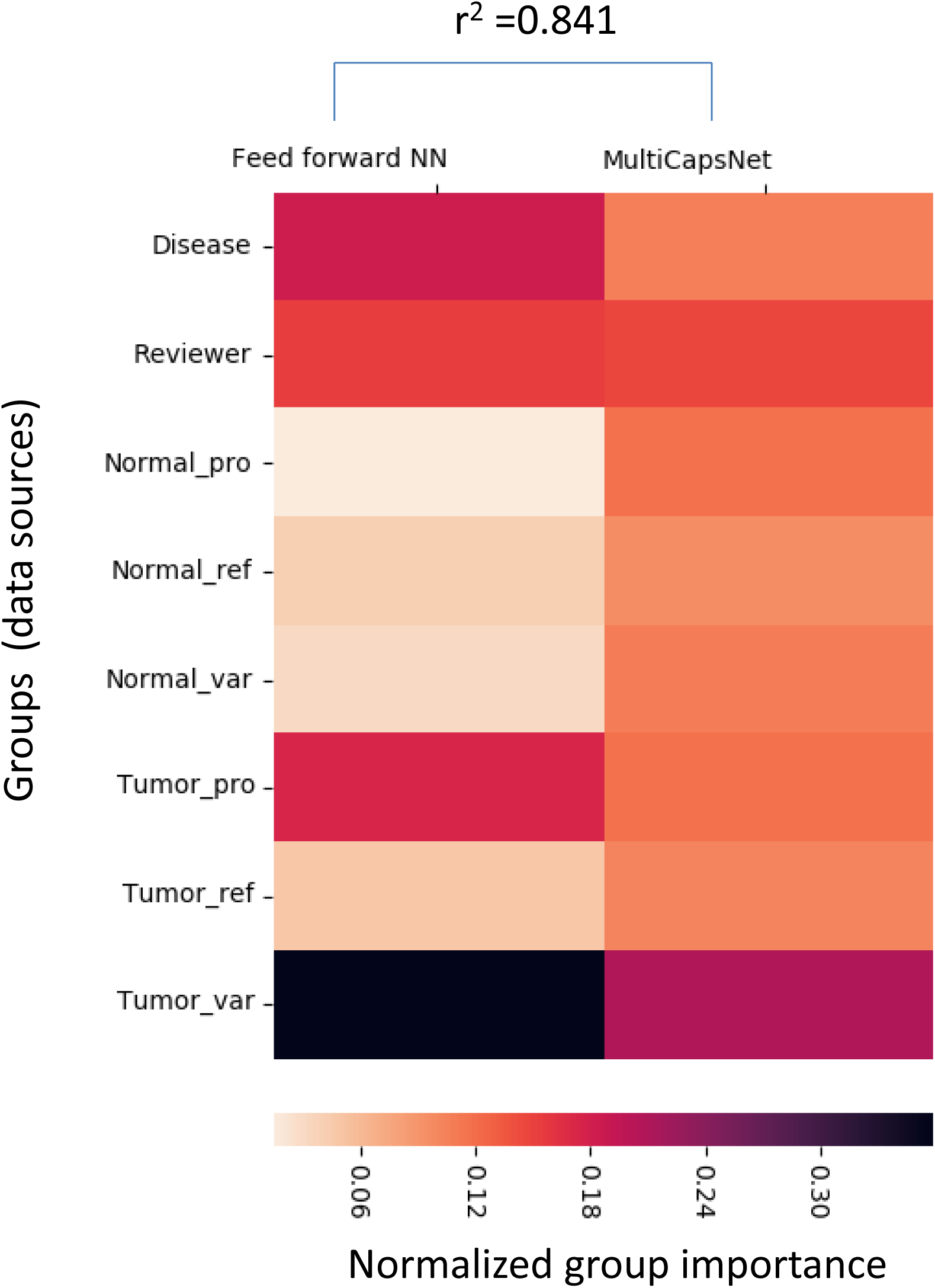
The normalized group (data source) importance. The MultiCapsNet group importance is highly correlated to the previous deep learning model group importance

**Fig. 4:**
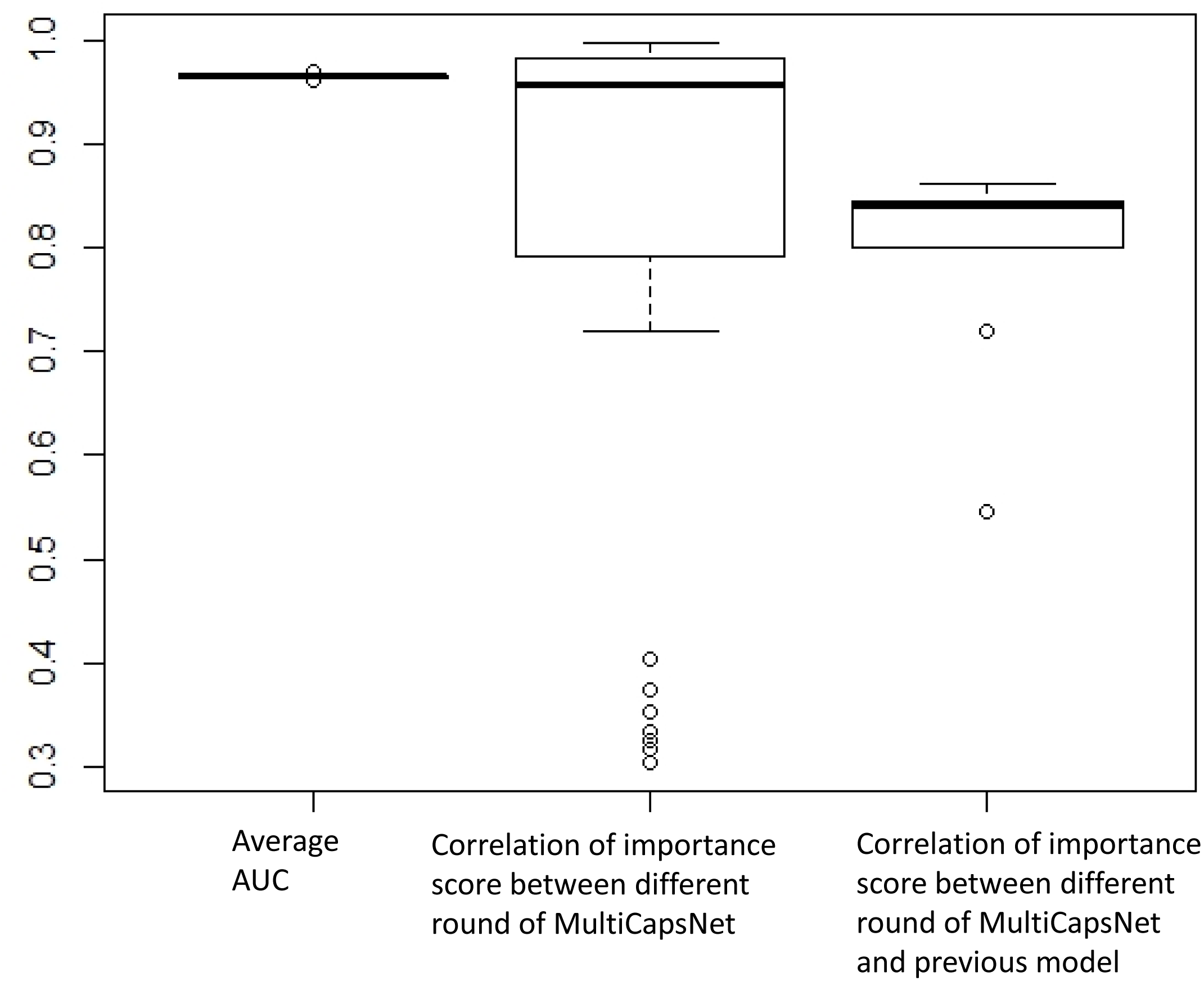
The consistency of MultiCapsNet model. The five lines from top to bottom in each boxplot represent upper hinge, upper quartile, median, lower quartile and lower hinge, respectively. The sample size of each column is 9, 36 and 9, respectively.

**Fig. 5:**
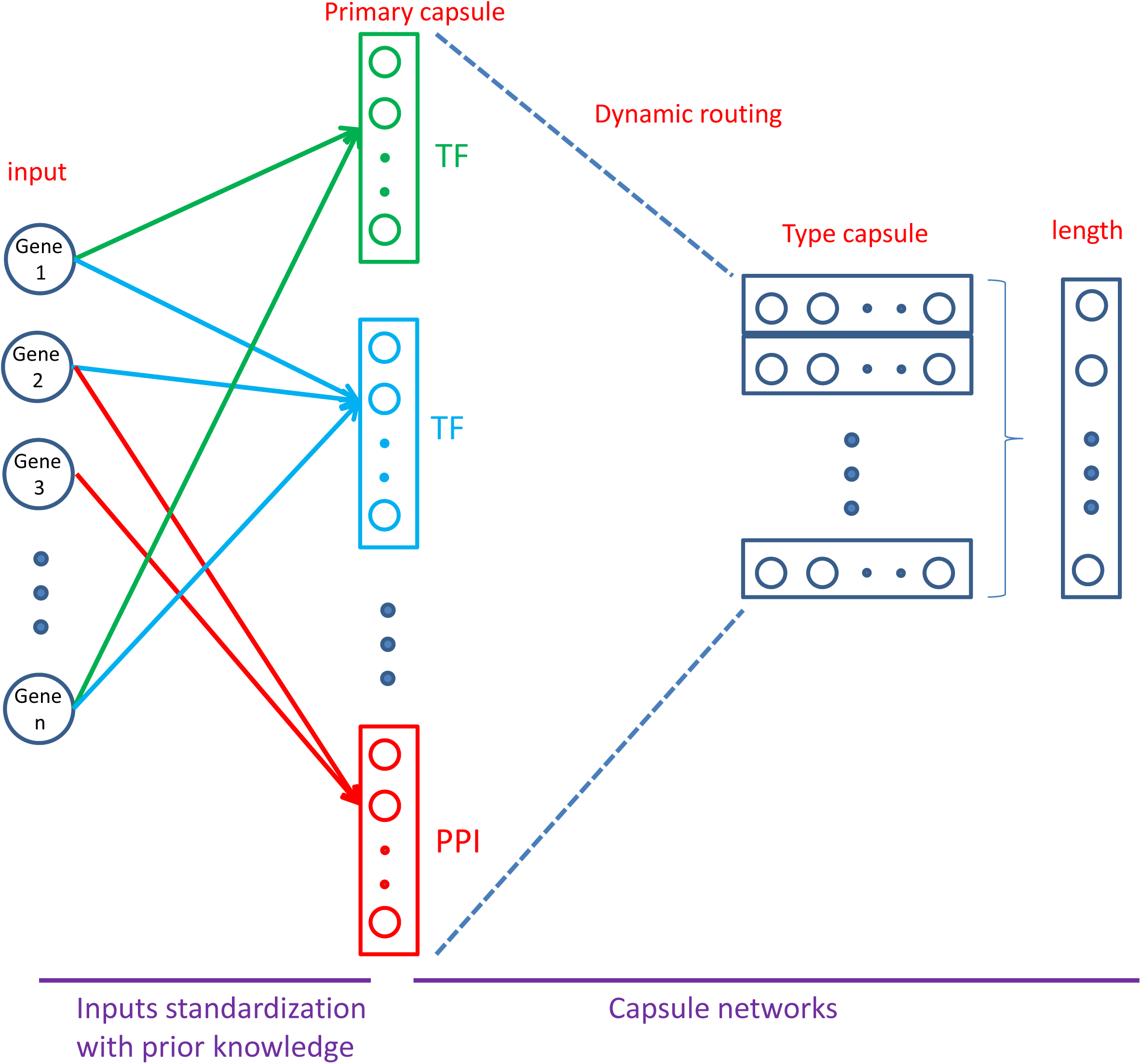
Architecture of MultiCapsNet integrates with prior knowledge. First layer consists of 696 parallel neural networks corresponding to 696 primary capsules labeled with either transcription factor (348) or protein-protein interaction cluster node (348). The inputs for each primary capsule include genes influenced by a transcription factor or involve in a protein-protein interaction network. The second layer is directly adopted from CapsNet for classification. The length of each label capsule represents the probability of input data belonging to the corresponding classification classes.

### Model with prior knowledge

The dataset, which is portion of mouse scRNA-seq data measured by microwell-seq, consists of over 4000 cells with 7 cell types and 9437 genes. The training accuracy reaches nearly 100% and the average validation accuracy is around 97%. The coupling coefficients of the mouse cell from training set with the same known cell type are grouped together and the average coupling coefficients are calculated for one specific cell type. The heatmaps visualizations of those average coupling coefficients are plotted (Fig. 6A). We also plot the overall heatmap, which incorporates the average coupling coefficients with type capsules corresponding to cell types from all training set (Fig. 6B). The average coupling coefficients could indicate how primary capsules contribute to cell type recognition. For each cell type, we rank the primary capsules according to their magnitude of average coupling coefficients. We use the labels of top ranked primary capsules, which are either transcription factor or PPI cluster nodes, to analyze how the transcription factor or PPI cluster nodes contribute to cell type recognition. The results show that the top ranked transcription factors or PPI cluster nodes often associate with the contributed cell type (Table 1). For example, the model reports GATA-binding factor 1 (GATA1) as a top contributor for dendritic cell recognition, and previous work indicates that Gata1 regulates dendritic-cell development and survival [23]. The Serum response factor (SRF) and Yin Yang 1 (YY1) are ranked as top contributors by the model for muscle cell recognition. As reported previously, SRF is required for skeletal muscle growth and maturation [24] and YY1 is associated with increased smooth muscle specific gene expression [25]. Nuclear receptor subfamily 3, group C, member 1(NR3C1) is ranked by model as a top contributor for cumulus cell recognition and is reported to be expressed by cumulus cell [26]. The model reports Forkhead box protein A2 (FOXA2) and Pre-B-cell leukemia transcription factor 1(PBX1) as top contributors for neuron recognition. And Foxa2 is reported to relate to A9 nigral dopamine neurons [27] and PBX1 transcriptional network is reported to control dopaminergic neuron development [28]. Homeobox protein aristaless-like 4 (ALX4) is ranked by model as a top contributor for cartilage cell recognition and it is reported to relevant with functions in the development of the craniofacial and/or appendicular skeleton [29]. We use the gene list enrichment analysis tool [30, 31] to examine the PPI cluster nodes. PPI_241 and PPI_199, both are high ranked contributors for AT2 cell, are reported to relate to lung with high probability by the enrichment analysis tool.

**Fig. 6:**
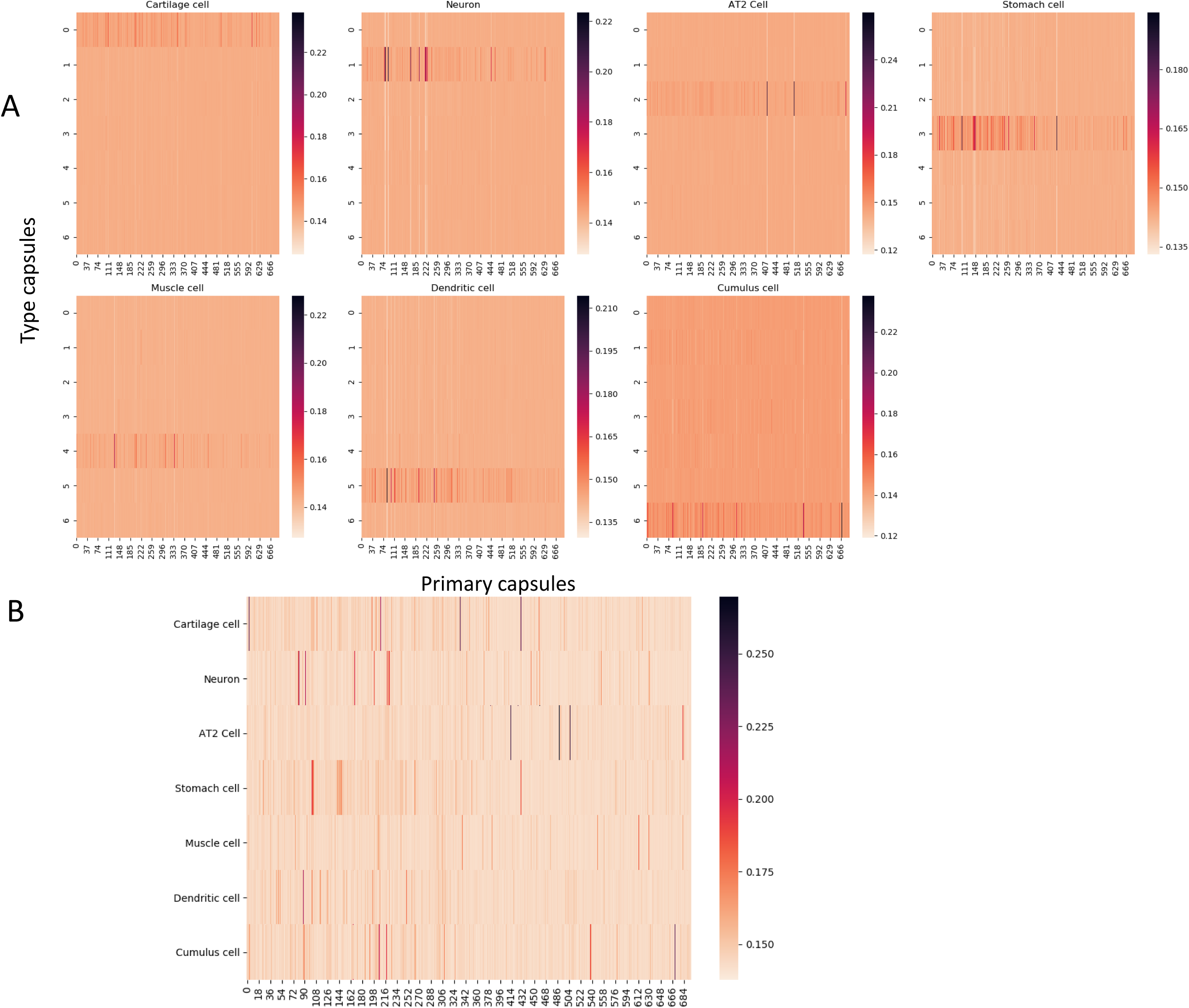
The heatmaps of coupling coefficients. **A.** Each heatmap represents average coupling coefficients of input cells from training set with one specific cell type that list on the top of each heatmap. The row represents type capsule and column represents primary capsule. The name of each type capsule could found in overall heatmap. **B.** The overall heatmap which combine the average coupling coefficients in the heatmaps of average coupling coefficients with type capsule is the same as the input cell type. The row represents primary capsules and column represents type capsules.

**Table 1:** Top ranked contributor (primary capsule) and their association to the cell type they contributed. The top ranked contributors given by the MuiltCapsNet model and the corresponding cell types they contribute are listed in the first and second columns. The third column indicates the average rank of the contributor. The score is calculated as following. Several training process are initiated with different training set-validation set division. The frequency of the contributors ranked in top 10, among those training process, is first counted, which present in the parentheses. Then, the average rank is calculated among those ranks list in the top 10. The reference and other information suggest an association between contributors and their contribute cells are list in the final column.

| Cell type | TF/PPI | Rank | reference/mouse gene atlas |
| --- | --- | --- | --- |
| Dendritic cell | GATA1 | 3(7/9) | Gata1 regulates dendritic-cell development and survival (23) |
| Muscle cell | SRF | 1(9/9) | Requirement for serum response factor for skeletal muscle growth and maturation (24) |
| Muscle cell | YY1 | 6(8/9) | Cytoplasmic YY1 is associated with increased smooth muscle-specific gene expression (25) |
| Cumulus cell | NR3C1 | 3(9/9) | Cumulus cells expressed genes for NR3C1 (26) |
| Cartilage cell | ALX4 | 2(9/9) | Alx4 is linked to the development of the craniofacial and/or appendicular skeleton (29) |
| Neuron | FOXA2 | 4(5/9) | Foxa2 related to A9 nigral dopamine neurons (27) |
| Neuron | PBX1 | 7(5/9) | PBX1 transcriptional network controls dopaminergic neuron development (28) |
| AT2 cell | ppi_241 | 3(9/9) | lung (Mouse Gene Atlas) |
| AT2 cell | ppi_199 | 3(9/9) | lung (Mouse Gene Atlas) |

## Discussion

In the first example, we demonstrate that the proposed MultiCapsNet model performs well in the variant call classification. Data sources with different data types, such as one-hot encoding and real value vectors, could be standardized into equal length vectors as primary capsules, and then pass the information into final layer capsules by dynamic routing. The importance of the data sources are measured by the sum of the overall average coupling coefficients as the co-product of the model training. These importance values are highly correlated with the importance values calculated by previous deep learning model, which is measured by average change in the AUC after randomly shuffling individual features.

In the second example, we incorporate PPI and PDI information into the structure of the MultiCapsNet model. This specified structure enforces the decoupling of the input, which is the scRNA-seq data, into several parts with each part corresponding to a set of genes influenced by a TF or involved in a network of protein-protein interaction. Each part of the decoupled input is viewed as a data source, and it connects to a primary capsule which is labeled as corresponding TF or PPI cluster nodes. Despite the number of the primary capsules is more than that of previous CapsNet model by one order of magnitude, the classification accuracy is very high. After training, the contribution of each primary capsule and its corresponding data source to the cell type recognition is revealed by the MultiCapsNet model as co-product of classification. The TFs or the PPI cluster nodes from the labels of the top ranked contributors are often relevant to the cell types they contributed.

To sum up, our MultiCapsNet model could integrate multiple input sources and standardize the inputs, then use the standardized information for classification through capsule network. In the variant call classification example, the data types are just limited to one-hot encoding and real value vectors. With suitable dataset, the MultiCapsNet could integrate and standardize more data types. For example, the sequence information could be converted into real-value vector by convolutional neural networks (CNN) [8], which could be included in the MultiCapsNet model as data sources. Otherwise, our MultiCapsNet model could also incorporate the prior knowledge through adjust the connection between layers according the specification of the prior knowledge. In scRNA-seq example, we just include PPI and PDI information. In the future, the information of complex, hierarchical structure of biological network would be introduced into MultiCapsNet model to better understand the intricacies of disease biology [6].

The MultiCapsNet model provides a framework for data integration, especially applicable in the context of multi-omics datasets with data from different sources and with different data types and formats, or in the scenario of the needs for prior knowledge. Once the data could be converted into read-value vectors through trainable parameters, then the data and the conversion process could be integrated into MultiCapsNet model as a building block. In this sense, MultiCapsNet model possesses enormous flexibility, and is applicable in many scenes, let alone that the importance of data sources could be measured accompanying the training step without any extra calculation step.

## Acknowledgments

This work was supported by grants from the National Key R&D Program of China [2018YFC0910402 to C.J.]; the National Natural Science Foundation of China [31571307 to C.J. and 61673070 to J.Z.]

## Supplementary Table

**Table 1.**
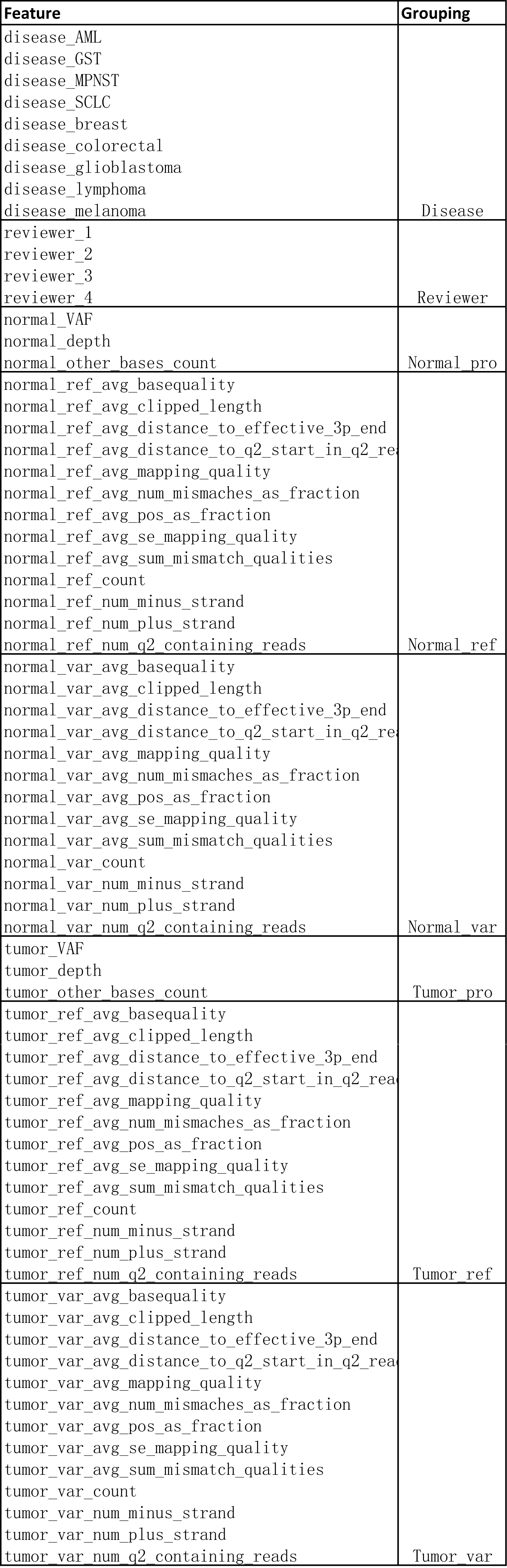
The 71 features and their grouping

## Supplementary Figure

**Fig. S1:**
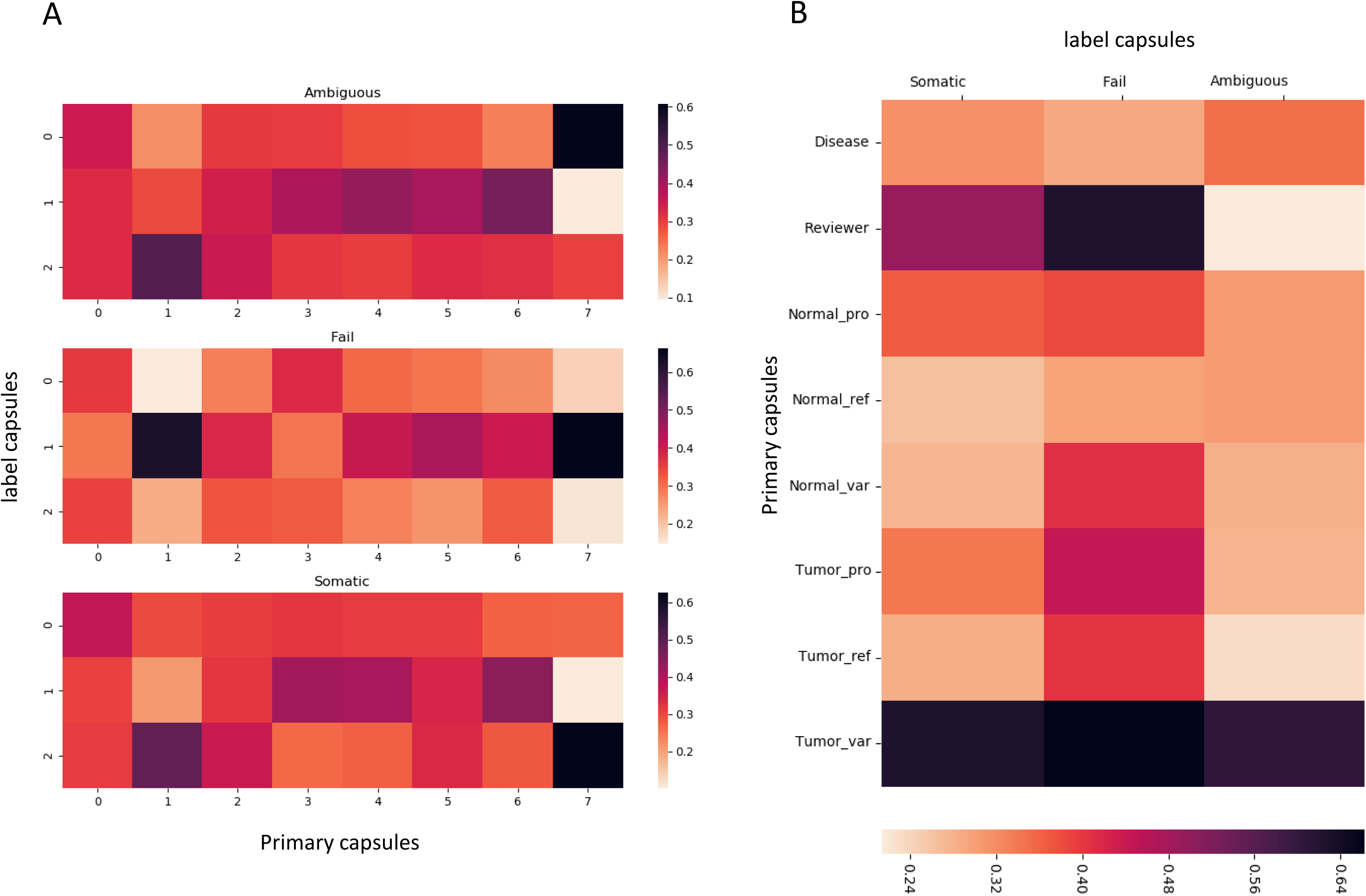
The heatmaps of coupling coefficients. **A.** Each heatmap represents average coupling coefficients of input variant calls with one specific label that list on the top of each heatmap. The row represents label capsule and column represents primary capsule. The name of each label capsule and primary capsule could found in overall heatmap. **B.** The overall heatmap which combine the average coupling coefficients in the heatmaps of average coupling coefficients with label capsule is the same as the input label. The row represents primary capsules and column represents label capsules.

